# Structural insight into *Pichia pastoris* fatty acid synthase

**DOI:** 10.1101/2021.01.27.428426

**Authors:** Joseph S. Snowden, Jehad Alzahrani, Lee Sherry, Martin Stacey, David J. Rowlands, Neil A. Ranson, Nicola J. Stonehouse

## Abstract

Type I fatty acid synthases (FASs) are critical metabolic enzymes which are common targets for bioengineering in the production of biofuels and other products. Serendipitously, we identified FAS as a contaminant in a cryoEM dataset of virus-like particles (VLPs) purified from *P. pastoris*, an important model organism and common expression system used in protein production. From these data, we determined the structure of *P. pastoris* FAS to 3.1 Å resolution. While the overall organisation of the complex was typical of type I FASs, we identified several differences in both structural and enzymatic domains through comparison with the prototypical yeast FAS from *S. cerevisiae*. Using focussed classification, we were also able to resolve and model the mobile acyl-carrier protein (ACP) domain, which is key for function. Ultimately, the structure reported here will be a useful resource for further efforts to engineer yeast FAS for synthesis of alternate products.

## Introduction

Fatty acid synthases (FASs) are critical metabolic enzymes for the endogenous biosynthesis of fatty acids in a diverse range of organisms. Through iterative cycles of chain elongation, FASs catalyse the synthesis of long-chain fatty acids that can produce raw materials for membrane bilayer synthesis, lipid anchors of peripheral membrane proteins, metabolic energy stores, or precursors for various fatty acid-derived signalling compounds^1^. In addition to their key physiological importance, microbial FAS systems are also a common target of metabolic engineering approaches, usually with the aim of generating short chain fatty acids for an expanded repertoire of fatty acid-derived chemicals, including chemicals with key industrial significance such as α-olefins^2–5^. A comprehensive structural and functional understanding of a diverse range of FAS systems is therefore paramount to the success of such complex rational engineering approaches.

Unlike the dissociated systems of individual enzymes that make up type II FAS systems in plants and bacteria^6^, Type I FAS systems in fungi and animals are large complexes that integrate the key enzymes required for fatty acid biosynthesis^7^,^8^. In particular, yeast type I FASs are 2.6 MDa hetero-dodecameric complexes with an α,_6_β_6_ configuration, which form a cage-like structure comprising two dome-shaped reaction chambers on either side of a central platform^9–13^. Along with structural support regions, enzymatic domains form the walls of the reaction chambers, with most active sites facing the chamber interior. The mobile acyl-carrier protein (ACP) domain tethers the growing fatty acid chain and carries it between active sites, so each can act on this in turn^13^,^14^. The phosphopantetheine transferase (PPT) domain, which catalyses the attachment of a phosphopantetheinyl group to the active site serine residue of the ACP (necessary for ACP activity), is also an integral part of yeast FAS complexes, and is located on the exterior of the cage^12^,^15^.

A deep body of work underlies our current structural understanding of yeast FAS, although FAS structure determination to date has been challenging. A cryoEM study of *S. cerevisiae* FAS revealed a number of differences with X-ray crystal structures of fungal FASs, suggesting that crystal contacts had altered the conformation of FAS from its unconstrained structure in solution^13^. A more recent attempt to determine the solution structure of *S. cerevisiae* FAS at a resolution sufficient for molecular modelling was hindered by partial denaturation induced by interactions with the air-water interface, an effect that was ameliorated by coating grids with hydrophilized graphene, ultimately leading to a 3.1-Å resolution structure when sample preparation was optimised^16^,^17^.

While our understanding of yeast FAS structure is growing, our library of structures remains incomplete. The vast majority of published solution structures of FAS are from *S. cerevisiae*^13^,^16–19^. To date, no structures are available for *Pichia pastoris* FAS from either cryoEM or X-ray crystallography. *P. pastoris* (*Komagataella spp*.) is a methylotrophic yeast which is used extensively for recombinant protein production^20^. In recent years, it has also been the target of metabolic engineering approaches^21^. *P. pastoris* has a major advantage over other yeast strains like *S. cerevisiae*, as it produces relatively little ethanol, making the maintenance of productive, high-cell density fermenter cultures more tractable^22^. It has also been suggested that *P. pastoris* cells, in contrast to cells of the plant *Arabidopsis* sp., are highly tolerant of free fatty acids^23^.

Here, we report the 3.1-Å resolution solution structure of *P. pastoris* FAS, determined by cryoEM. We initially identified FAS as a contaminant in a *P. pastoris-derived* virus-like particle (VLP) preparation being used for structural studies, and processed these data to yield detailed structural information on FAS, including density for the mobile ACP domain. Comparison of our atomic model with that of *S. cerevisiae* revealed interesting structural differences throughout the complex within both structural and enzymatic domains. This included movement of the malonyl/palmitoyl transferase domain at the level of domain packing, and alternative rotamers in the catalytic centres of the acyl transferase and enoylreductase domains.

## Results

### CryoEM structure determination of *Pichia pastoris* fatty acid synthase

Virus-like particles (VLPs) composed of hepatitis B virus tandem core proteins^24^ with a SUMO-binding affimer inserted into the major immunodominant region (MIR)^25^ were expressed in transformed *P. pastoris* cells for structural and vaccine studies (construct available upon request). Expression was induced with methanol and the VLPs purified from the yeast lysate by sucrose gradient centrifugation. Image analysis of cryoEM data revealed, in addition to the VLPs, non-VLP-like particles which were recognised as yeast FAS complexes due to their characteristic morphology – two hollow chambers on either side of a central platform. Further inspection of individual micrographs revealed that FAS particles were well-separated from VLPs, indicating that FAS was not co-purified as a result of an association with the VLPs (Fig 1A).

**Figure 1.**
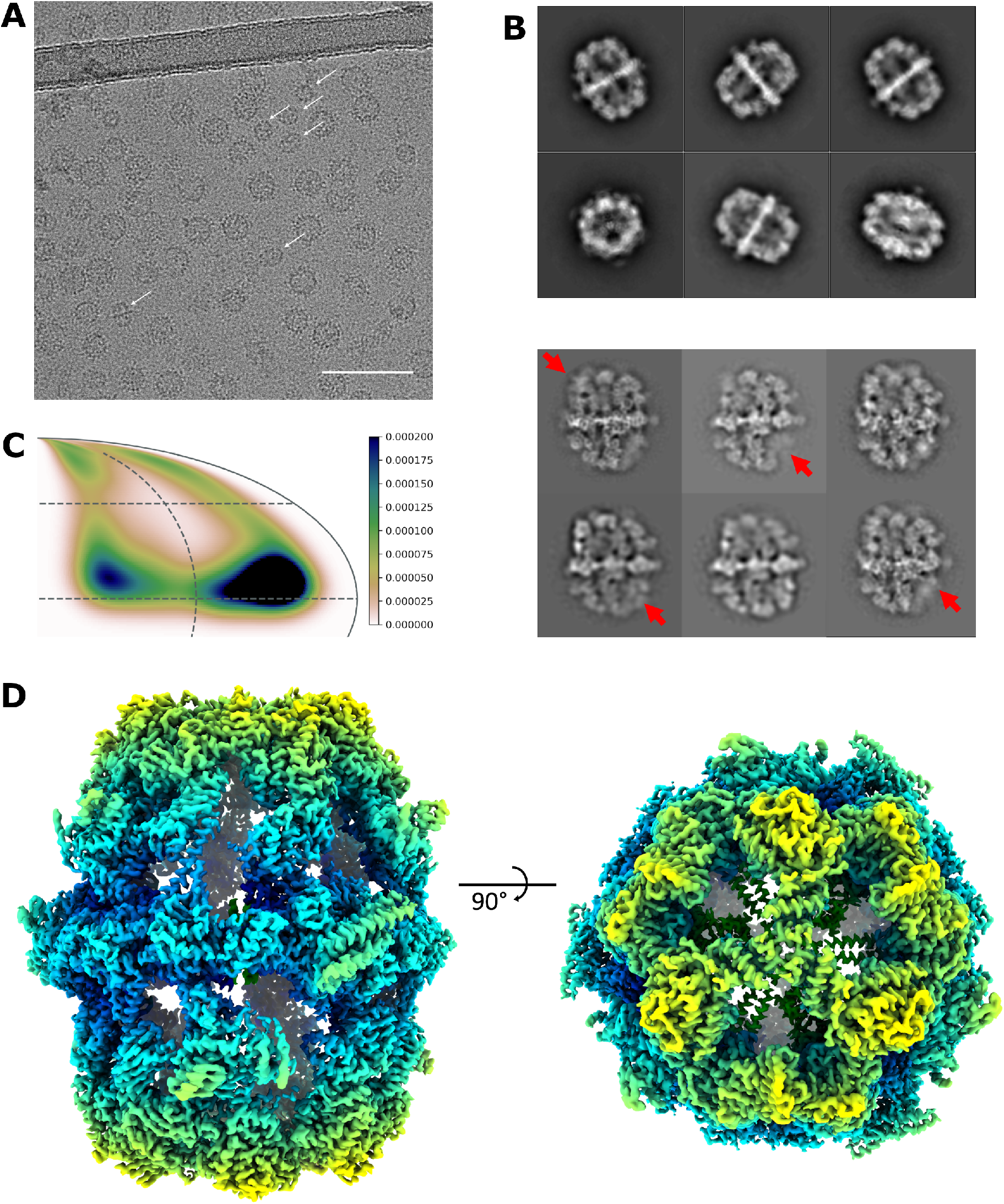
Determination of P. pastoris FAS structure by cryoEM. **(A)** Raw micrograph showing VLPs with contaminating FAS particles. FAS particles are indicated by white arrows. Scale bar shows 100 nm. **(B)** The most populated classes from 2D classification of P. pastoris FAS (after excluding ‘junk’ classes of FAS over thick carbon support) (upper) compared with 2D class averages of S. cerevisiae FAS showing partial denaturation (indicated by red arrows) from D’Imprima et al. (2019)^16^ (licensed under <u>CC BY 4.0</u>) (lower). **(C)** Orientation distribution map of particles contributing to the final reconstruction of the FAS asymmetric unit. The map shows the probability that any particle was assigned the orientation indicated, according to the scale shown. **(D)** CryoEM density map of P. pastoris FAS, coloured radially.

In contrast to a recent study that reported significant air-water interface-induced denaturation of FAS particles by cryoEM (which could be solved with deposition of hydrophilized graphene on the grid surface)^16^, we saw no signs of denaturation in our data set, judged by 2D class averages (Fig 1B). While some particles were discarded after the initial 2D classification, ~67% of these were actually VLPs or areas of carbon film that were inadvertently selected during autopicking, and the remaining discarded classes mainly comprised apparently intact FAS particles over thick carbon. It is therefore difficult to estimate how many particles were excluded because of damage that was not visible in class averages. There was evidence of some preferential orientation effects (perhaps induced by adsorption to the carbon film), but this was not a significant problem, as a reasonable coverage of potential orientations within the asymmetric unit of the D3 symmetric structure (i.e., a single α and β subunit) was achieved (Fig 1C).

Classes corresponding to FAS were taken forward and processed separately, yielding a 3.1 Å reconstruction from ~37,000 particles, with parts of the central platform resolved to ~2.7 Å (Fig 1D, Supplementary Fig S1). In comparison, ~28,000 particles of *S. cerevisiae* FAS were used to generate a 4.0-Å resolution structure^16^. More recently, ~15,000 particles of *S. cerevisiae* FAS led to a 3.1-Å resolution map, although in this case the FAS was incubated with NADPH and malonyl-CoA before vitrification in order to drive synthesis of fatty acids to completion, increasing particle homogeneity^17^.

### The 3.1-Å resolution solution structure of *P. pastoris* FAS

The overall structure of the *P. pastoris* FAS reported here was typical of a yeast FAS. Density was present for most domains of the α- and β-subunits, including acyl transferase (AT), ketoacyl reductase (KR), ketoacyl synthase (KS), enoylreductase (ER), dehydratase (DH) and malonyl/palmitoyl transferase (MPT) domains (Fig 2A,B). ACP density was weak (only becoming fully apparent when inspecting the map at a threshold of approx. 1 σ) and poorly resolved, as was also the case for previous EM density maps of *S. cerevisiae* FAS (e.g., EMD-10420, EMD-0178). However, the approximate position of the ACP could be identified within the reaction chamber. There was no density apparent for the PPT region, as discussed further below.

**Figure 2.**
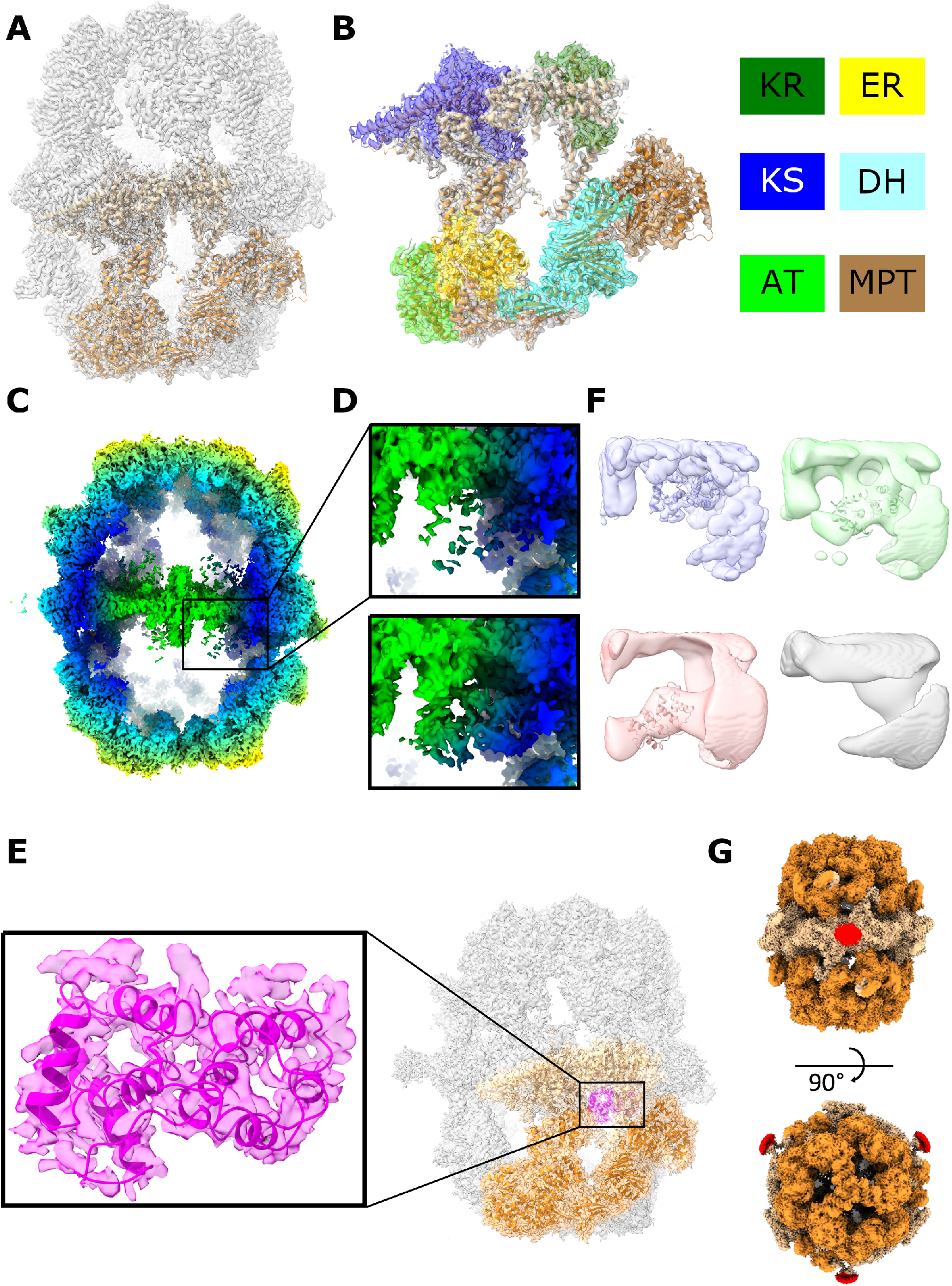
The molecular structure of P. pastoris FAS. **(A)** Atomic coordinates for the asymmetric unit of P. pastoris FAS, comprising α (light orange) and β (dark orange) subunits, with the density map overlaid. **(B)** Enlarged view of the asymmetric unit, highlighting different enzymatic domains according to the colour scheme indicated. Grey-coloured regions are non-enzymatic, structural domains. Domain boundaries were assigned based on equivalent domain boundaries from S. cerevisiae FAS. KR – ketoacyl reductase, KS – ketoacyl synthase, AT – acetyl transferase, ER – enoylreductase, DH – dehydratase, MPT – malonyl/palmitoyl transferase. **(C)** Clipped view of the sharpened FAS density map, coloured according to radius. The box highlights the region of the reaction chamber selected for the first focussed 3D classification. **(D)** Enlarged image of the region depicted by the box in (C) (upper) compared with the sharpened, asymmetric reconstruction of FAS derived from an ACP density-containing class from the first focussed 3D classification (lower). All classes from the first focussed classification are given in Supplementary Fig S2A. **(E)** Atomic coordinates for the ACP domain of FAS fitted into ACP density from the map shown in (D, bottom). ACP is also shown in the context of the full FAS complex. **(F)** Representative classes from the second focussed 3D classification (with expanded masked region) of the complex interior. Atomic coordinates for ACP are shown fitted into the three classes containing putative ACP density to give an indication of size. One class shows no putative density for ACP (lower right). All maps are shown at the same contour level. All classes from the second focussed classification are given in Supplementary Fig S2B. **(G)** Density map of FAS shown at a low contour level (i.e., including weak density), coloured according to subunit (α - light orange, β - dark orange). Unexplained density, not accounted for by FAS subunits α or β, is highlighted in red.

The map was sufficiently resolved to allow us to build an atomic model of *P. pastoris* FAS. An initial homology model for the asymmetric unit was generated using SWISS-MODEL^26^ and rigid-body fitted into the density map. The fit of model to density was visually inspected and corrected, before the model was symmetrised and refined to improve its fit to the density and geometry. The a subunit is mostly resolved, though several segments of the peptide sequence (including the ACP and PPT domains) are missing (residues 96-322, 535-598 and 1752-1879). The β subunit is practically complete, lacking only a small number of residues at the N- and C-termini (residues 1-9 and 2064-2069). We also observed density for a ligand bound within the active site of the ER domain (for reference, see Fig 3G), presumably a molecule of FMN.

**Figure 3.**
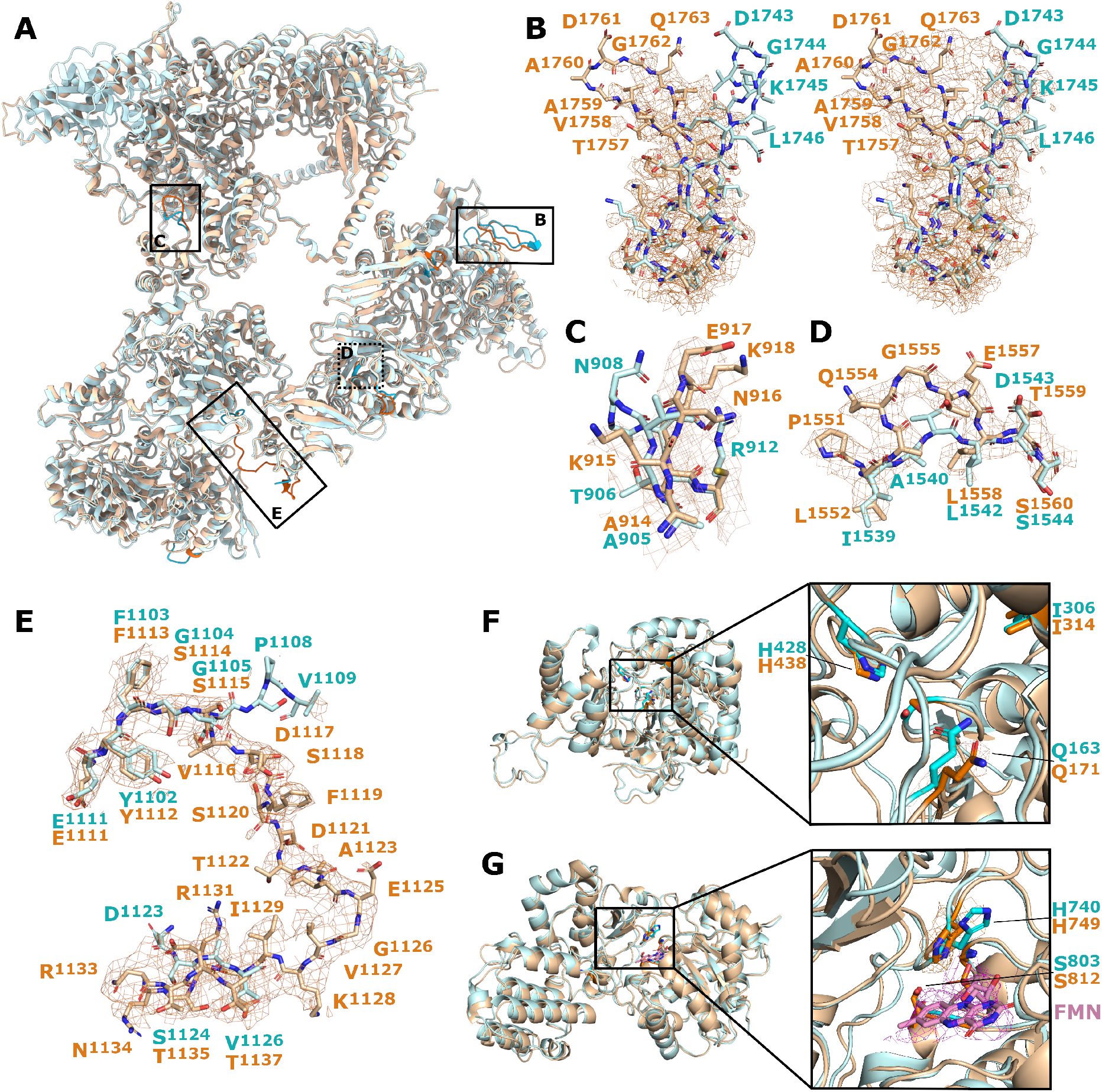
Unique structural features of P. pastoris FAS. **(A)** Overlay of P. pastoris FAS (orange) and S. cerevisiae FAS (PDB 6TA1, blue) from Joppe et al. (2020)^17^, with differences highlighted by stronger colours. Enlarged images are indicated on the map with boxes. **(B)** Loop region comprising residues 1753-1768, highlighting the different positioning compared to the equivalent region from S. cerevisiae FAS. The P. pastoris density map is overlaid at high contour level (left) and low contour level (right). **(C-E)** Other regions from different parts of the protein are highlighted and compared with equivalent regions of S. cerevisiae FAS, including the loop comprising residues 914-919 **(C)**, the beta strand comprising residues 1553-1558 **(D)** and the additional resolved linking region comprising residues 1113-1135 **(E)**. **(F)** The AT domain, with enlarged image highlighting different positioning of the active site residue Q171. **(G)** The ER domain, with enlarged image highlighting different positioning of the active site residue H749. FMN and its corresponding density are shown in pink.

In an attempt to improve the quality of the cryoEM density corresponding to the ACP domain, we performed focussed 3D classification within the reaction chamber of the complex (Fig 2C). While classification with a regularisation parameter (‘T’ number) of 20 did not reveal any meaningful classes, using a higher value of 40 (i.e., putting a higher relative weighting on the experimental data than on the prior/reference) yielded a class containing relatively well resolved ACP density (Supplementary Fig S2A). Particles from this class were taken forward for asymmetric reconstruction, yielding a complete FAS map of 3.1-Å resolution (Fig 2D). In the latter map, the additional density is consistent with the ACP domain of the a subunit of FAS, given the fit-to-density of an ACP domain homology model. This additional density was largely missing in the original map. Ultimately, the new density was of sufficient quality to build an atomic model of the ACP domain (Fig 2E). Interestingly, ACP domain residue S178 (which tethers the growing fatty acid chain) is located ~18 Å away from the KS active site, which is in agreement with the length of the PPT arm (18 Å)^27^. Some classes contained additional low-resolution density at the edge of the masked region used for focussed classification (Supplementary Fig S2A). To examine whether this corresponded to ACP domains positioned differently within the complex interior that were not fully contained within the masked region, focussed classification was repeated with an expanded mask (Supplementary Fig S2B). In addition to a class containing well-resolved ACP density, multiple classes containing low-resolution ACP-sized density in different positions were observed (Fig 2F). However, these classes comprised relatively few subparticles and asymmetric reconstruction only resulted in poor resolution (~8 Å).

Focussed classification was also performed in an attempt to resolve the PPT domain. While a low-resolution class was identified as containing PPT density (Supplementary Fig S3), further reconstruction did not yield density sufficient for model building.

Interestingly, we observed additional density on the exterior of the FAS complex, not accounted for by the atomic model (Fig 2G). This density is weaker than for the rest of the complex and is not well resolved, indicating that the component is flexible and/or that there is not full occupancy. The additional, external density appears to extend outwards from V534 (a subunit). While residues 535-598 are not resolved, the density remains localised close to V534 and does not provide a logical connecting trace to A599. The extra density was also present in a 4.3-Å resolution reconstruction determined from the same data but without the imposition of symmetry, suggesting that this is not simply an artefact of symmetry averaging. However, given the poor resolution of this region of the map, we were unable to determine its identity.

### Distinct structural features of *P. pastoris* FAS

To identify unique structural features of *P. pastoris* FAS, we carried out a comparative structural analysis. While PDBeFold^28^ suggested that the closest structural matches to *P. pastoris* FAS were cryoEM structures of *C. albicans* FAS (PDB 6U5W, PDB 6U5V)^18^, we chose to compare our structure with *S. cerevisiae* FAS, a better characterised FAS with greater relevance to metabolic engineering attempts. At the peptide sequence level, *P. pastoris* FAS α and β subunits show 69% and 63% identity with their equivalents in *S. cerevisiae*. We compared our structure with a recently published 3.1-Å resolution structure of FAS from *S. cerevisiae* (PDB 6TA1)^17^. After aligning the two structures, a root mean-square deviation (RMSD) value between equivalent Ca atom pairs was calculated as 7.1 Å, indicating a notable level of difference between the two models (Fig 3A). However, aligning the models by individual subunits gave RMSD values of 1.7 Å (α-α), and 2.2 Å (β-β), suggesting that this may be partially attributable to differences in the spatial relationship between the α and β subunits in each model. Visual inspection confirmed that several domains appeared shifted, particularly the MPT domain.

Closer examination revealed many differences in the path traced by the peptide backbone. One instance of this is a loop in the MPT domain, which is shifted by 16 Å at its turning point (D1761) (Fig 3B). Interestingly, when displaying density at a low threshold so that even weak density is visible (0.5 σ), we also saw density tracing the alternative loop conformation seen in the *S. cerevisiae* yeast structure. Other loops and linker regions throughout the β subunit also showed obvious differences in conformation, including in the DH domain (F1283-V1289, E1505-I1511), MPT domain (A1859-Y1870), and structural domains/linker regions (G75-N78, A914-G919) (Fig 3C). Furthermore, a small sequence insertion of three amino acids introduced a kink into the outer strand of a β-sheet within the DH domain (A1553-L1558) (Fig 3D). Of particular interest is a sequence of residues (F1113-T1135) linking the ER domain to the DH-adjacent structural domain, which was mostly unmodeled in the *S. cerevisiae* FAS structure (Fig 3E). For the structure reported here, this linker region was well-resolved and differed significantly from the terminal parts of the linker that are resolved in PDB 6TA1.

There are also subtle rotameric differences in some of the enzymatic domain active sites. In particular, Q171 was shifted within the active site of the AT domain (Fig 3F). H749 of the ER domain was also pointing in the opposite direction to the equivalent histidine in *S. cerevisiae* FAS (Fig 3G).

## Discussion

We serendipitously identified fatty acid synthase, which is constitutively expressed in yeast^29^, as a contaminant in a cryoEM data set of a hepatitis B core VLP preparation purified from *P. pastoris*. Subsequent analysis of raw micrographs revealed that FAS was readily separated from the VLPs. As such, we think it unlikely that FAS co-purified as a result of a specific interaction with the VLPs. Rather, its presence in the VLP preparation was probably as a result of its mass being sufficiently similar to co-purify on sucrose density gradients (FAS 2.6 MDa; *T*=3 VLP 4.7 MDa; *T*=4 VLP 6.3 MDa). By processing this ‘contaminant’ data and using focussed 3D classification techniques, we were able to determine the structure of the *P. pastoris* FAS to 3.1 Å resolution, and build an atomic model for the complex, including a model for the mobile ACP domain.

Interestingly, we did not observe signs of partial denaturation of FAS in 2D class averages, as was seen previously for a different yeast FAS structure^16^. It is possible that the presence of VLPs in the sample affected the ice thickness or crowded the air-water interface (which is known to induce denaturation^16^), reducing any interactions between the interface and FAS. However, we suspect that the lack of partial denaturation is more likely a result of the continuous thin carbon film used for making these cryoEM grids, which may trap FAS particles away from the air-water interface. While the presence of a thin carbon film adds a background that reduces the signal-to-noise ratio of particles (and can therefore limit the resolution), we were able to resolve our map to 3.1 Å, equivalent to that of the structure reported by Joppe *et al*. (2020)^17^ and sufficient to build a molecular model for FAS. Therefore, using a continuous thin carbon film may be a simple alternative to coating grids with hydrophilized graphene, when the size of the target protein (and hence its signal strength) is similar to FAS.

The overall structure was typical of a yeast FAS. As has previously been observed for yeast FAS structures, the PPT domain was missing^16^ and the quality of the ACP domain density was relatively poor. In this case, density corresponding to the ACP domain was initially of insufficient quality for molecular modelling, but this was resolved by focussed classification. Given the known mobile nature of the ACP domain^13^, this was unsurprising. Focussed classification led to an improvement in the strength and quality of the ACP density that was sufficient to allow atomic modelling, probably as result of grouping together only subparticles with ACP domains in the same relative position (adjacent to the KS domain, as has been observed in several other yeast FAS structures^17^,^27^) at the time of vitrification. Use of an expanded mask for focussed classification revealed classes with putative ACP density in other positions within the complex interior. These classes contained relatively few subparticles, suggesting that the vast majority of ACP domains were adjacent to the KS domain, with the remainder either missing or occupying other positions. The lack of PPT domain density is likely to be a result of loss of this domain from most complexes during sample purification/preparation, as has previously been noted^16^,^17^. Since our initial aim was to purify VLPs, not FAS, no special care was taken to ensure the complex was kept intact (although the ER domain cofactor FMN was clearly present in the density map, providing evidence that the complex was still likely to have been active). While focussed classification was not sufficient to resolve the PPT domain, one class did show weak density corresponding to the PPT domain, suggesting that it was retained in at least some particles.

We identified a number of structural differences between the *P. pastoris* FAS structure reported here and a recent 3.1-Å resolution structure from *S. cerevisiae* (PDB 6TA1)^17^, another member of the yeast family *Saccharomycetaceae*. The MPT domain was distinctly shifted relative to its position in *S. cerevisiae* FAS. This is of particular importance, because this domain is the target of one approach to engineer FAS complexes to produce short chain fatty acids^4^. Many intradomain differences affected structural (non-enzymatic) components of the complex, including a loop within the MPT domain from S1752 – K1768. This loop was particularly interesting, as very weak density tracing the backbone of the alternative (*S. cerevisiae*) conformation also became visible at low contour thresholds. This is indicative of a mixed population of conformations in *P. pastoris* FAS, with most complexes sampling the conformation modelled, and a smaller subpopulation sampling the conformation seen for *S. cerevisiae* FAS. This loop may plausibly be dynamic, an idea reinforced by the local resolution of the loop being poorer than average, at around 3.7 – 4.1 Å resolution (Supplementary Fig S1).

We also identified differences in the active sites of two enzymatic domains. In the AT domain, Q171 forms part of a tight hydrogen bond network around the active serine (S282) and its backbone amide also contributes to an oxyanion hole, which plays a role in promoting the transfer of malonyl to the thiol group of the ACP^6^,^12^. While the alternative rotamer observed here is unlikely to affect the oxyanion hole, the structure suggests a subtle alteration in the arrangement/spacing of the hydrogen bond network at the catalytic centre of the AT domain. Aside from Q171, the catalytic histidine of the ER domain (H749) was seen to be in a ‘flipped’ conformation relative to its equivalent in *S. cerevisiae* FAS^12^,^17^. Interestingly, the conformation reported here is much closer to that seen in a crystal structure of *S. pneumoniae* FabK in its active state (PDB 2Z6I), a homologous enzyme that forms part of its type II FAS system^30^. Whether these changes to active site residues have functional significance is unclear. However, it should be noted that the *S. cerevisiae* FAS was incubated with NADPH and malonyl-CoA prior to vitrification in order to drive FA synthesis to completion^17^. The *P. pastoris* FAS reported here received no such treatment and as such, reaction status was likely less uniform and potentially incomplete.

In conclusion, despite non-optimised sample preparation and using just ~37,000 particles, we were able to determine the structure of *P. pastoris* FAS to 3.1-Å resolution and identify structural differences with a related yeast FAS. As well as highlighting the importance of considering ‘contaminant data’ in cryoEM data sets, the structure reported here could prove a useful resource for future efforts to engineer yeast FAS.

## Methods

### Cells

FAS was purified from PichiaPink™ yeast strain one (Invitrogen, USA), which was grown according to the manufacturer’s instructions.

### Yeast transformation and induction

While the focus of this work was FAS, which is constitutively expressed in yeast^29^, our initial aim was to use yeast to express and purify VLPs composed of hepatitis B virus tandem core proteins^24^ with a SUMO-binding affimer^25^ inserted into the major immunodominant region (MIR) (construct available upon request) for structural characterisation. As such, VLP expression was induced in the transformed yeast prior to purification, as described previously^31^. Briefly, glycerol stocks known to exhibit good levels of expression were cultured in yeast extract-peptone-dextrose (YPD), supplemented with 50 μg/mL ampicillin, at 28°C, 250 rpm for 48 h. 2 ml of the high-density starter culture was then transferred to 200 mL YPD and incubated for another 24 h under the same conditions. To induce expression of VLPs, cells were pelleted (1500 × *g*, 20 min) and resuspended in 200 mL yeast extract-peptone-methanol (YPM; 0.5% [v/v] methanol) for a further 48-hour incubation, with methanol added to a final concentration of 0.5% (v/v) 24 h post-induction. Cells were subsequently collected by pelleting at 2000 × *g* (20 min), resuspended in 30 mL breaking buffer (50 mM sodium phosphate, 1 mM phenylmethylsulfonyl fluoride [PMSF], 1 mM EDTA, 5% glycerol, pH 7.4) and stored at −20°C prior to purification.

### Co-purification of FAS with VLPs

Though our aim was to purify VLPs (as described previously^31^) for structural characterisation, FAS co-purified with the particles. Frozen *P. pastoris* stocks that had been induced to express VLPs were thawed and incubated with 0.1% (v/v) Triton-X100 for 30 min. Cells were lysed using a CF-1 cell disruptor at ~275 MPa and cooled to 4°C, before the lysate was centrifuged first at 4000 rpm (30 min, 4°C), then at 10,000 × *g* (30 min, 4°C) to remove insoluble material. The supernatant was supplemented with 2 mM MgCl and treated with HS-nuclease (2BScientific) for 1.5 h at room temperature, then mixed with saturated ammonium sulphate solution (40% [v/v]) and incubated overnight at 4°C for precipitation. Precipitated protein was recovered by pelleting at 4000 rpm (30 min, 4°C) and solubilised by resuspending in PBS (supplemented with 0.1% [v/v] Triton-X100). After further centrifugation at 4000 rpm (30 min, 4°C), then 10,000 × *g* (30 min, 4°C) to remove any remaining insoluble material, the supernatant was pelleted through a 30% sucrose cushion (w/v) at 151,000 × *g* (3.5 h, 4°C, SW-32 Ti). The pellet was resuspended in PBS (supplemented with 1% [v/v] NP-40 and 0.5% [w/v] sodium deoxycholate), then clarified at 10,000 × *g* (10 min, 4°C) before loading onto a 15-45% (w/v) sucrose gradient for ultracentrifugation at 151,000 × *g* (3 h, 4°C). 2 ml fractions were collected and those containing VLPs were identified by western blot. A centrifugal filter (100K, 5 mL, Pall Life Sciences) was used according to the manufacturer’s instructions to remove sucrose and concentrate the peak fraction to 0.5 mL.

### Electron microscopy

CryoEM grids were prepared by applying 3 μL of a VLP preparation with contaminating FAS complex in PBS to lacey carbon 400-mesh copper grids coated with a <3-nm continuous carbon film (Agar Scientific, Stansted, UK), after glow-discharging the grids in air (10 mA, 30 seconds) using a PELCO easiGlow™ Glow Discharge System (Ted Pella). Following application of sample, grids were incubated at 8°C, 80% relative humidity for 30 seconds, then blotted to remove excess liquid and vitrified in liquid nitrogen-cooled liquid ethane using a LEICA EM GP plunge freezing device (Leica Microsystems, Wetzlar, Germany). Blotting time parameters were varied to increase the likelihood of recovering a grid with optimal ice thickness. Following plunge freezing, grids were stored in liquid nitrogen. Grids were imaged using an FEI Titan Krios transmission electron microscope (ABSL, University of Leeds, Leeds, UK) operating at 300 kV. A magnification of 75,000× was used for a calibrated object sampling of 1.065 Å/pixel. Detailed information on data collection parameters is provided in Table S1.

### Image processing

The RELION 3.0 pipeline^32^ was used for image processing. Raw micrographs were first motion corrected using MotionCor2^33^, then CTF parameters were estimated using Gctf^34^. Particles were selected from a subset of micrographs by manual picking and were then differentiated by 2D classification. High quality 2D classes were used as templates for autopicking the complete data set. A low picking threshold was specified to autopicking to minimize the number of ‘missed’ VLPs. Picked particles were 2× down-sampled, then subject to 2D classification (with CTFs ignored until the first peak), leading to the identification of FAS-containing classes. 2D classes containing FAS were taken forward for initial model generation and used as templates for FAS-specific autopicking. Autopicked FAS particles were extracted without down-sampling and subjected to two rounds of 2D classification (first with CTFs ignored until the first peak, then without). Particles in high quality classes were taken forward for 3D refinement (with D3 symmetry applied) with subsequent masking and sharpening. Following this, several cycles of CTF refinement (with per-particle astigmatism correction and beamtilt estimation), Bayesian polishing and 3D refinement (with masking and use of solvent-flattened FSCs) were performed to improve the resolution of the map. After sharpening, the resolution of the final map was determined using the ‘gold standard’ Fourier shell correlation criterion (FSC = 0.143) (Table S1, Figure S1). RELION was used to estimate local resolution and generate a local resolution-filtered map.

To resolve the flexible ACP, focussed 3D classification was performed as described previously^35–37^. Briefly, a cylindrical mask was generated in SPIDER^38^ and resampled onto the D3-symmetric density map of FAS, such that it was positioned over the weak ACP density. The relion_symmetry_expand tool was used to assign 6 symmetrically redundant orientations to each particle contributing to a symmetric reconstruction, such that the ACP domain in each symmetry-related position would be included in the classification. These symmetry-expanded particles were then subject to masked 3D classification without re-alignment using a regularisation parameter (“T” number) of 40. Particles contributing to the ACP density-containing class were taken forward for asymmetric reconstruction using the relion_reconstruct tool.

### Model building and refinement

To generate a preliminary model for the asymmetric unit of FAS, the peptide sequences of *P. pastoris* FAS α (NCBI: XP_002490414.1) and β (NCBI: XP_002489642.1) subunits were used to generate a homology model using the SWISS-MODEL web server^26^. The homology model was rigid-body fitted into the sharpened FAS density map using UCSF Chimera^39^, then refined in Coot^40^. The refined model was symmetrised in UCSF Chimera, then subject to real space refinement in Phenix^41^ to improve simultaneously the fit of the model to the density map and model geometry. Several iterations between refinement in Coot and refinement in Phenix were performed before validating the model with Molprobity^42^.

### Visualization and structure comparison

Density map and atomic model visualization was performed and figures were generated using UCSF Chimera^39^ and UCSF Chimera X^43^. Structural alignment and RMSD calculations were performed using the ‘MatchMaker’ tool with default settings in UCSF Chimera. PDBeFold (the protein structure comparison service at EBI [http://www.ebi.ac.uk/msd-srv/ssm])28 was used to probe structural similarity between the FAS structure reported here and other molecular structures within the PDB. Particle orientation distribution was visualized using an adapted version^44^ of a script from Naydenova & Russo (2017)^45^.

### Quantification and statistical analysis

Where appropriate, statistical details are given in the Method Details section. Table S1 contains quantitative parameters related to data collection and image processing. Table S2 contains validation statistics related to model building.

## Data Availability Statement

The *P. pastoris* FAS cryoEM map was deposited in the Electron Microscopy Data Bank (EMD-12138) along with the reconstructed ACP domain-containing map from focussed classification (EMD-12139). The atomic coordinates for the FAS asymmetric unit (PDB-7BC4) and the ACP domain (PDB-7BC5) were deposited in the Protein Data Bank.

## Acknowledgments

We are grateful to the Wellcome Trust for PhD Studentship support to JSS (102174/B/13/Z) and to the Saudi Ministry of Education for studentship support to JA. LS is funded by the WHO (2019/883397-O, “Generation of virus free polio vaccine – phase IV”). Electron microscopy was performed in the Astbury Biostructure Laboratory, which was funded by the University of Leeds and the Wellcome Trust (108466/Z/15/Z). We also wish to thank David Klebl (University of Leeds) for assistance with identifying FAS, and we are very grateful to Glyn Hemsworth (University of Leeds) and Alison Baker (University of Leeds) for critical reading of the manuscript.

## Author Contributions

JA and LS maintained *P. pastoris;* JA purified FAS; JSS prepared samples for cryoEM, carried out cryoEM data collection and performed image processing; JSS, JA, DJR, NAR and NJS analysed data; JSS, NAR and NJS wrote the paper; LS, MS, DJR, NAR and NJS provided supervision.

## Additional Information

The authors declare no competing interests.

