## Supplementary Information for "Structural insight into *Pichia pastoris* fatty acid synthase"

**Supplementary Table S1 (related to Figure 1).** Quantitative parameters related to cryo-EM data collection.

| | **FAS** |
|---|---|
| Microscope | FEI Titan Krios |
| Camera | Falcon III |
| Voltage (kV) | 300 |
| Pixel size (Å) | 1.065 |
| Total dose ($e^-/Å^2$) | 60.1 |
| Number of frames | 40 |
| Defocus range (µm) | –0.8 to –3.0 |
| Number of micrographs | 3643 |
| Acquisition software | Thermo Scientific EPU |
| Motion correction | MotionCor2 |
| CTF estimation | GCTF |
| Image processing | Relion 3.0 |
| Particles contributed | 37,054 |
| B-factor | -112 |
| Resolution (FSC=0.143) (Å) | 3.1 |
| Map resolution range (Å) | 2.7 – 4.9 |

**Supplementary Table S2 (related to Figures 2 and 3).** Quantitative parameters and validation statistics related to atomic model building.

| | FAS | FAS ACP domain |
|---|---|---|
| **PDB ID** | 7BC4 | 7BC5 |
| **Residues modelled** | α: 1-95, 323-534, 599-1751; β: 10 - 2063 | α: 139 - 299 |
| **RMSD** | | |
| *Bond lengths (Å)* | 0.0082 | 0.0055 |
| *Bond angles (°)* | 1.20 | 1.32 |
| **Validation** | | |
| *All-atom clashscore* | 5.38 | 7.83 |
| *Molprobity score* | 1.77 | 1.92 |
| *Rotamer outliers (%)* | 1.19 | 0.76 |
| **Ramachandran plot** | | |
| *Favoured (%)* | 93.61 | 91.82 |
| *Allowed (%)* | 6.39 | 8.18 |
| *Outliers (%)* | 0.0 | 0.0 |

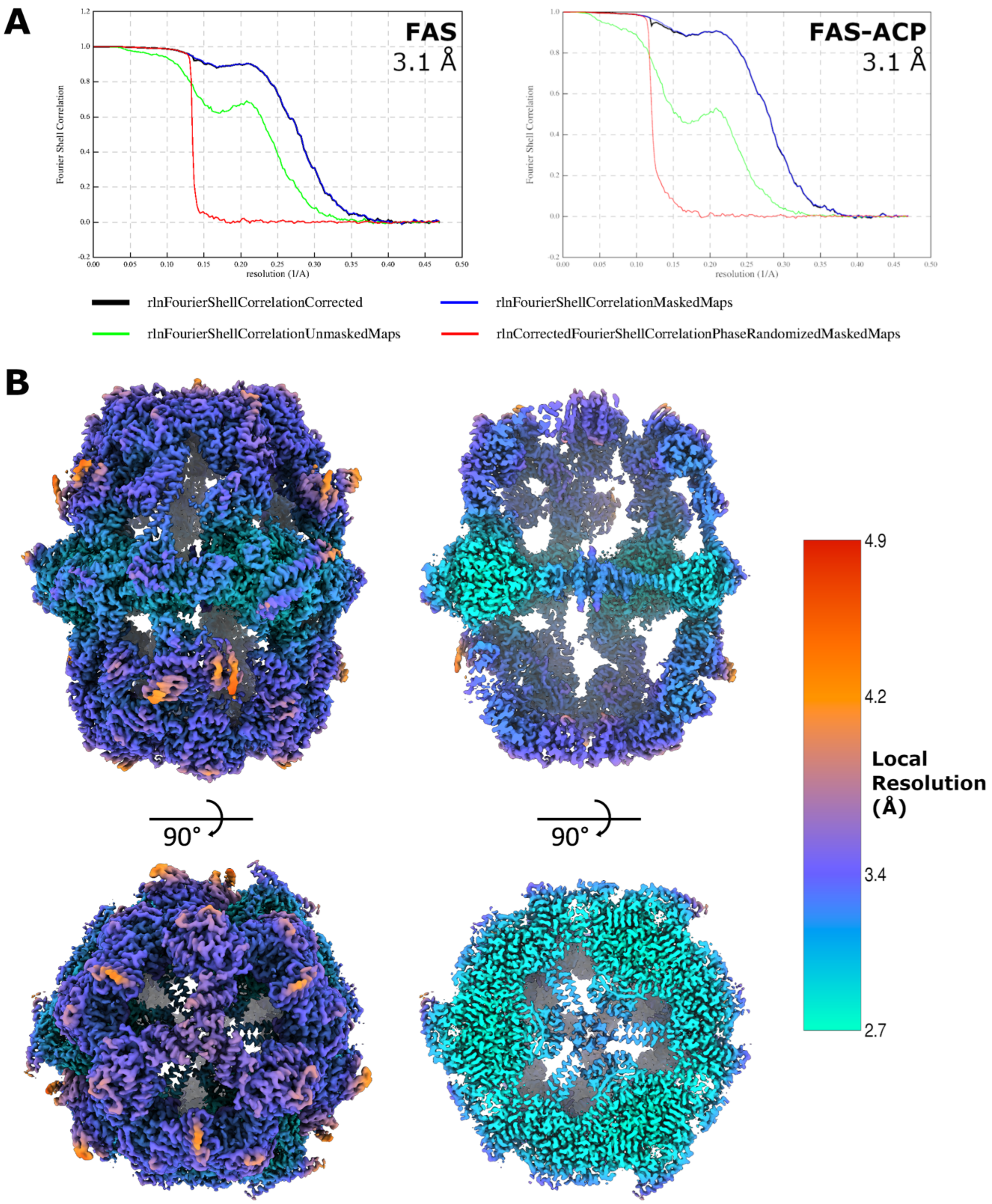


**Supplementary Figure S1 (related to Figures 1 and 2).** The 3.1-Å resolution reconstruction of *P. pastoris* FAS. **(A)** Fourier shell correlation (FSC) plots for the full symmetric FAS reconstruction (FAS, left) and the reconstruction of FAS from the focussed class containing improved ACP density (FAS-ACP, right). The resolution for each map was determined using the FSC=0.143 criterion. **(B)** Isosurface representation of the FAS reconstruction, filtered by local resolution and coloured according to the local resolution scale indicated.

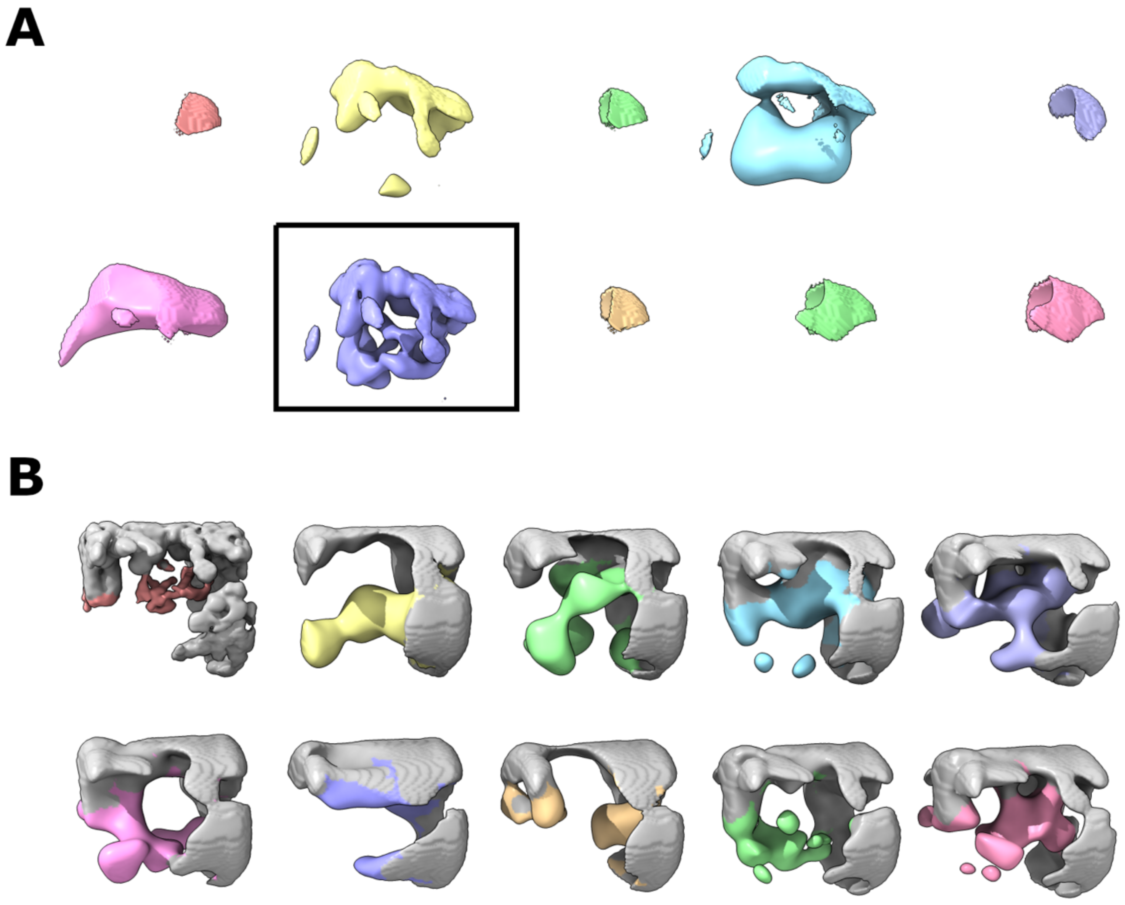


**Supplementary Figure S2 (related to Figure 2). (A)** All classes from focussed 3D classification of the ACP-containing region of the reaction chamber. All classes are shown at the same contour level. Particles from the boxed class were used for asymmetric reconstruction of FAS. **(B)** All classes from focussed 3D classification of the ACP-containing region of the reaction chamber using an expanded masked region. Grey regions correspond to density within 5.5 Å of FAS atomic coordinates without the ACP domain (i.e., the outer wall and central platform that surround the ACP domain-containing interior chamber). All classes are shown at the same contour level. Several classes are also shown in Fig 2F.

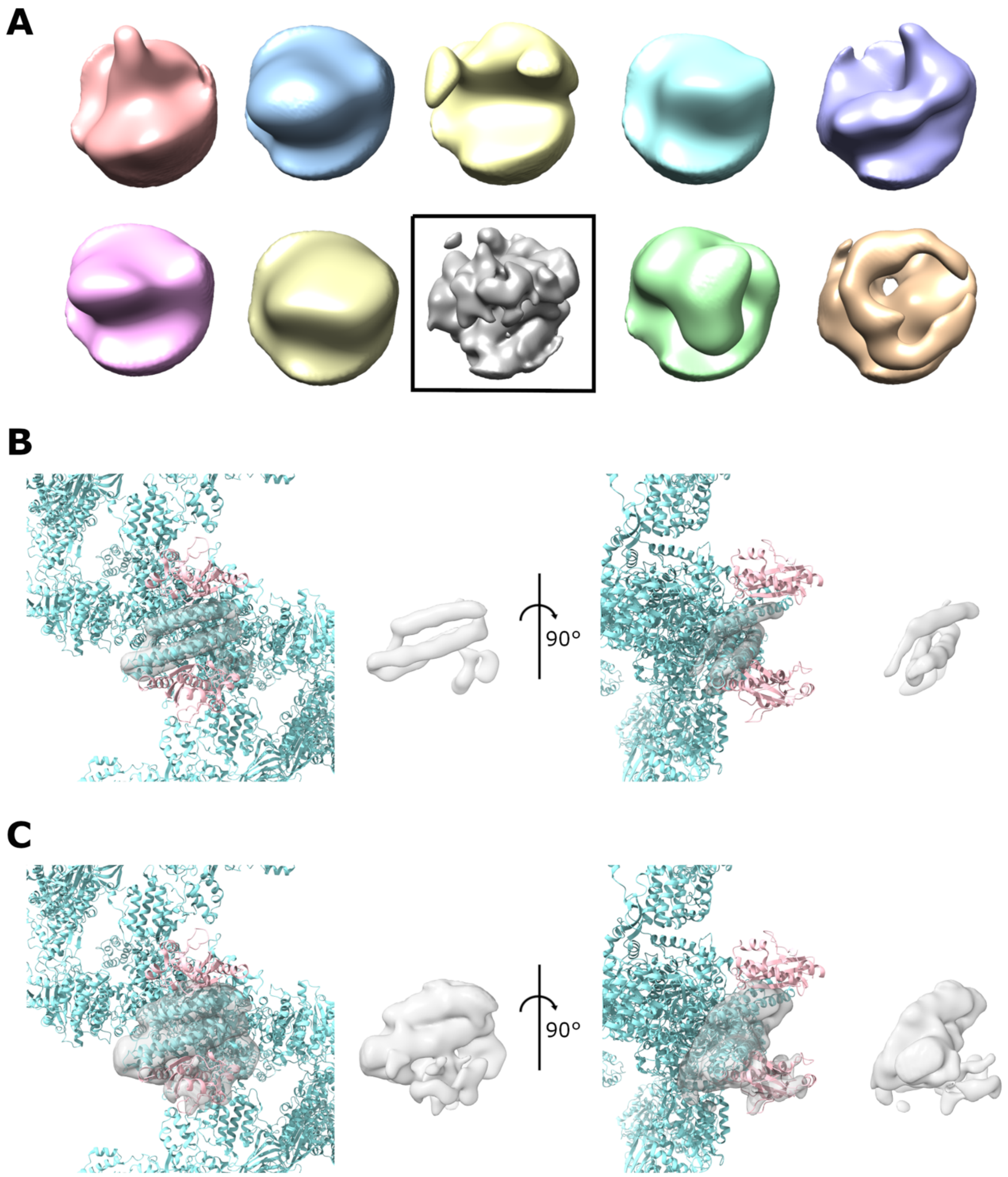


**Supplementary Figure S3 (related to Figure 2).** Focussed classification of the PPT domain. **(A)** All classes from focussed 3D classification of the region expected to contain PPT domain density. **(B,C)** PPT density-containing class (indicated by the box in (A)) shown at a high contour threshold (only strong density shown) **(B)** and a low contour threshold (weak density also visible) **(C).** Density is shown from two different viewing angles, either alone or overlaid with the model of *S. cerevisiae* FAS (PDB 6TA1) which contains atomic coordinates for the PPT domain (highlighted in pink).
